# Pyramids and cascades: a synthesis of food chain functioning and stability

**DOI:** 10.1101/361246

**Authors:** Matthieu Barbier, Michel Loreau

## Abstract

Food chain theory is one of the cornerstones of ecology, providing many of its basic predictions, such as biomass pyramids, trophic cascades and predator-prey oscillations. Yet, ninety years into this theory, the conditions under which these patterns may occur and persist in nature remain subject to debate. Rather than address each pattern in isolation, we propose that they must be understood together, calling for synthesis in a fragmented landscape of theoretical and empirical results. As a first step, we propose a minimal theory that combines the long-standing energetic and dynamical approaches of food chains. We chart theoretical predictions on a concise map, where two main regimes emerge: across various functioning and stability metrics, one regime is characterized by pyramidal patterns, the other by cascade patterns. The axes of this map combine key physiological and ecological variables, such as metabolic rates and self-regulation. A quantitative comparison with data sheds light on conflicting theoretical predictions and empirical puzzles, from size spectra to causes of trophic cascade strength. We conclude that drawing systematic connections between various existing approaches to food chains, and between their predictions on functioning and stability, is a crucial step in confronting this theory to real ecosystems.

## INTRODUCTION

The concept of food chain, since its formulation by Elton [1], has become one of the most widely studied in empirical and theoretical ecology [2]. It provides a lens through which we can understand many ecological phenomena, by partitioning species into trophic levels, describing how biomass or energy is distributed among these levels, and studying their dynamics, from population cycles to trophic cascades [3].

Yet, its fundamental predictions have a checkered history of success outside of textbook examples. When predicted patterns are not observed, it is often unclear whether the issue lies with the theory and its application, or the influence of other factors such as complex interactions and spatial fluxes [4–11]. Even when predictions *do* agree with empirical data, they might not be sufficient to ascertain the underlying trophic structure [12–15].

This lack of consensus may be due to an increasing fragmentation of the literature. Food chains are characterized by an interconnected set of structural, functional and dynamical properties. Yet, these properties have grown into separate topics of investigation, influenced by two main paradigms (Fig. 1).

**FIG. 1.**
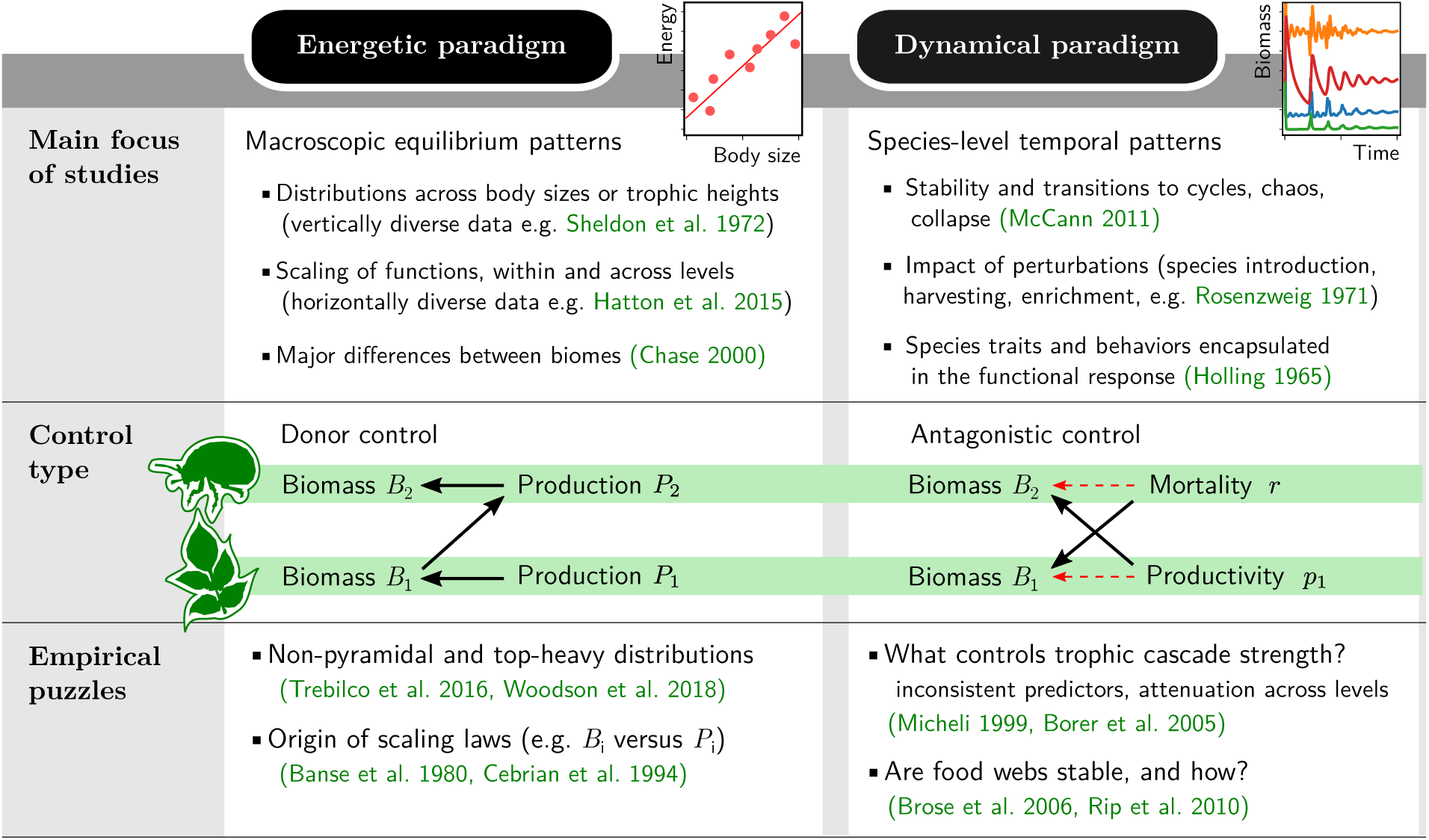
The energetic and dynamical approaches to food chain behaviors. The *energetic* paradigm focuses on static and macroscopic patterns, while the *dynamical* paradigm emphasizes temporal patterns and individual species characteristics. Beyond these complementary concerns, each paradigm also offers a different vision of how trophic levels interact. In donor control, each level’s biomass scales with its production, fixed by the biomass of the level below. We explain in the main text that this scaling requires self-regulation. Antagonistic control, seen in dynamical models without self-regulation (8), is less intuitive: each level’s biomass is set either by its prey’s productivity, or by its predator’s mortality, and is uncorrelated with its own productivity. These two assumptions lead to clashing theoretical expectations, and empirical puzzles specific to each approach. We list here four such challenges that we address in the main text.

The *energetic* paradigm, following Hutchinson and Lindeman [16], focuses on macroscopic energy flows up the food chain. Based on scaling arguments and statistical relationships, it is associated with large-scale empirical and applied studies, notably in marine systems [17–19]. It seeks laws describing how aggregated metrics of functioning (e.g. biomass, growth or consumption) vary across trophic levels or body sizes [20, 21], how they scale with each other within a level [22], and how they relate to environmental or physiological parameters. These static relationships are built upon the implicit assumption that ecosystems operate at, or close to, some steady regime.

The *dynamical* paradigm, following Lotka [23] and Volterra [24], focuses on how the impacts of predation ripple down the food chain. It often operates at the scale of individual species, with an attention to the response and behavior of predators and prey. Drawing on a rich mathematical literature, it emphasizes the importance of feedback mechanisms and nonequilibrium phenomena such as cycles or collapse [25]. Some of its predictions, including trophic cascades and the paradox of enrichment [26], have prompted extensive empirical investigations, but theory and empirics have rarely been matched at a quantitative level [27].

While these two paradigms usually address different questions, it is possible to apply one’s methods to the other’s objects, revealing unexpected conflicts. Jonsson [28] noted that energetic arguments produce pyramidal biomass distributions, whereas dynamical models often do not. We will exhibit a number of other discrepancies in relations between metabolism, biomass, productivity and stability.

Yet, we believe that these theoretical approaches can and must be embedded within a single framework. This systematic viewpoint has been argued for at a conceptual level by Leibold *et al*. [29]. But the field is still lacking a general quantitative formalism: a concise map of all essential food chain behaviors and their interplay, along with the key ecological parameters that govern them.

As a first step toward this synthesis, we propose an approach that captures aspects of both paradigms. We use a simple dynamical food chain model that includes metabolic rates and self-regulating mechanisms within a trophic level. In distinct parameter ranges, this model can recover either the energetic paradigm’s scaling relationships, or the stability patterns investigated in the dynamical paradigm.

We show that the modelling assumptions of the two paradigms can be understood as opposite corners in a multidimensional spectrum (Fig. 2), and we systematically explore how functioning and stability properties vary across the full spectrum. The most important axis is the relative strength of predation and self-regulation, which predicts the transition between two qualitatively different regimes. In the bottom-up regime dominated by self-regulation, all the properties studied here, from biomass to variability, exhibit pyramidal patterns. In the top-down regime dominated by predation, these properties all display cascade (alternating) patterns.

**FIG. 2.**
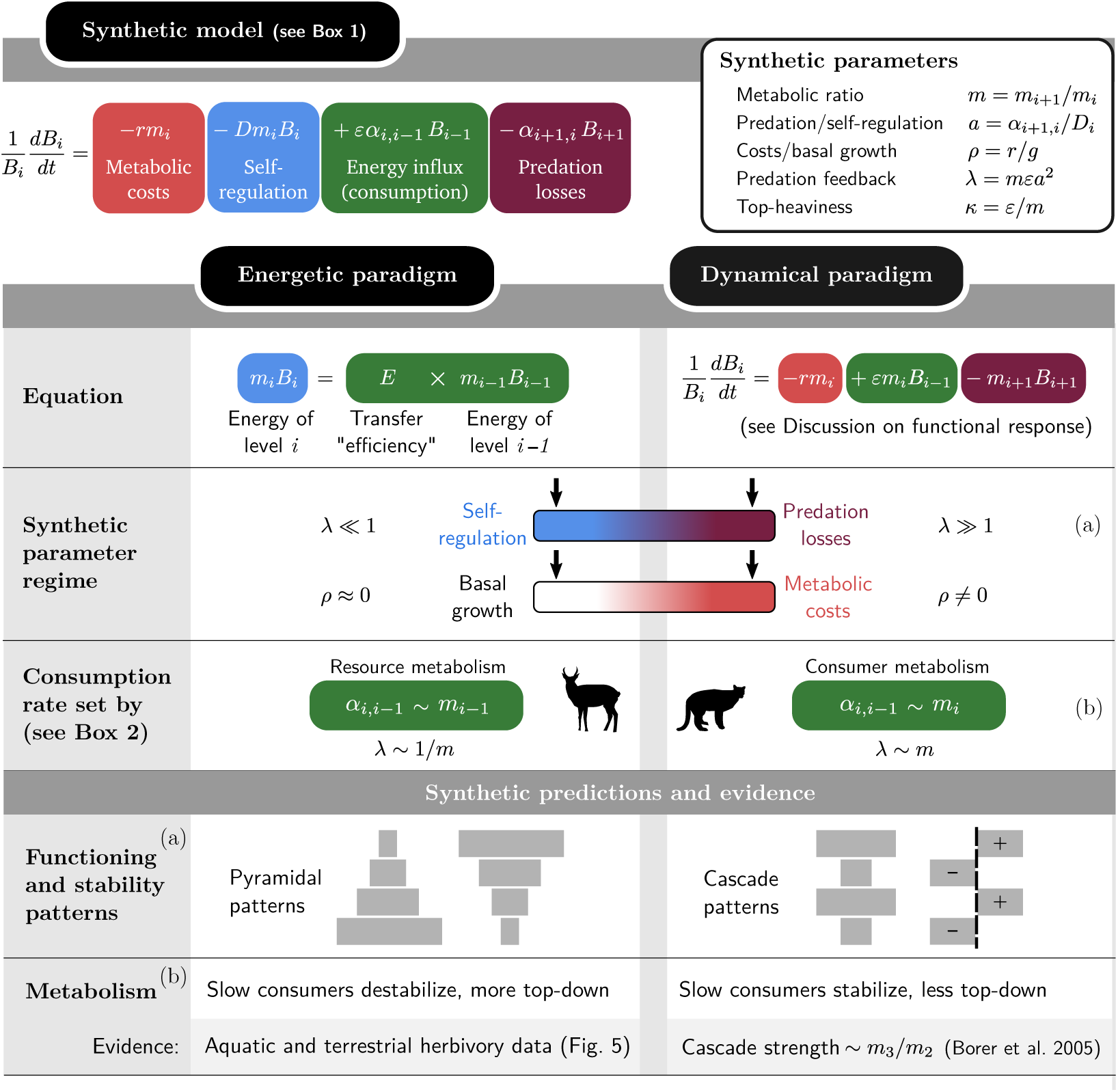
Synthesizing the energetic and dynamical paradigms and deriving systematic predictions. In the *energetic* paradigm, the biomass *B*_*i*_ of trophic level *i* is often predicted using the heuristic formula (7) reproduced in the figure. The *dynamical* paradigm emphasizes the role of predators in regulating their prey, and explores the role of different functional responses [26] such as the Lotka-Volterra model (8) shown here. By embedding both paradigms in our synthetic model (defined in Box 1), we can highlight their differences. At equilibrium, *dB*_*i*_/*dt* = 0, we can identify the equations term-by-term. Terms are color-coded by their dependence in *B*_*i*_, *B*_*i*−1_ and *B*_*i*+1_. Each approach emphasizes some terms over others, as symbolized by the color gradients and arrows in “Synthetic parameter regime”. We see that the energetic formula (7) neglects predation mortality (*λ* 1) and ignores metabolic losses (*ρ ≈* 0, equivalent to high basal growth). This leads to pyramidal patterns in various properties, from biomass (Fig. 3) to stability (Fig. 4). On the other hand, the dynamical paradigm emphasizes predation loss but ignores consumer self-regulation (right-hand arrows), leading to *cascade* patterns, i.e. alternating patterns of positive and negative, or high and low, values across trophic levels. In addition, the energetic formula assumes that consumption is proportional to resource metabolism, while dynamical models [30] generally assume it is proportional to consumer metabolism (Box 2). This leads to divergent predictions on the role of metabolism in biomass distribution and stability, see main text and Fig. 5. The labels (a) and (b) relate assumptions to predictions.

**FIG. 3.**
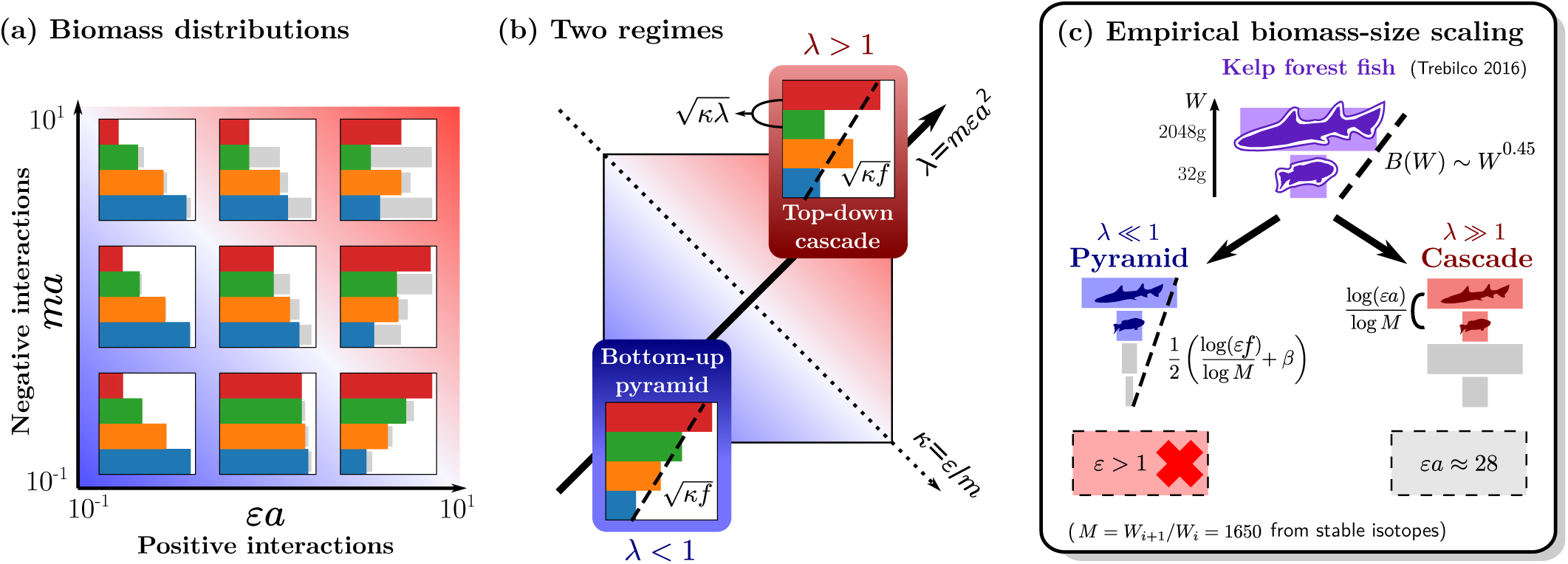
Theoretical map of equilibrium biomass patterns in a food chain, and application to an empirical puzzle. The parameters (defined in Box 1) are predator-prey metabolic ratio *m*, conversion efficiency *ε* and the strength of interactions *a* relative to self-interaction. Results are illustrated for a four-level chain with negligible metabolic losses *ρ* ≪ 1 (other examples in Supporting Information). **(a)** Biomass distribution. For each trophic level, a colored bar represents its equilibrium biomass (in log scale), and a grey bar represents the amount of biomass lost to its predator, reflecting the intensity of top-down control. **(b)** Two parameter regimes define the shape of the biomass distribution. The main diagonal *λ* = *mεa*^2^ distinguishes the region of bottom-up control *λ* < 1 with pyramids (both regular and inverted), from the region of top-down control *λ* > 1 with cascade (alternating) patterns. The other diagonal *κ* = *ε*/*m* affects the top-heaviness of the biomass distribution, as the global slope of the distribution is given by 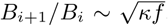. The fraction *f* of production lost to consumers is given by (14). In cascade patterns, low-biomass and high-biomass levels *i* and *i* + 1 alternate, with 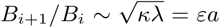 (larger than the global slope, which still holds betwen levels *B*_*i*+1_ and *B*_*i*−1_). **(c)** Identifying trophic structure from data. Given the biomass per trophic level and the mass ratio *M* between levels, we can compute the total biomass in each body size class *W*. Fish in kelp forests exhibit a top-heavy (positive) scaling *B*(*W*) *∼ W*^0.45^ [43]. We show in main text that this scaling cannot hold across multiple trophic levels, except for an unphysical value of conversion efficiency *ε* > 1 in formula (16). It may, however, be found between two adjacent levels under strong top-down control (17).

**FIG. 4.**
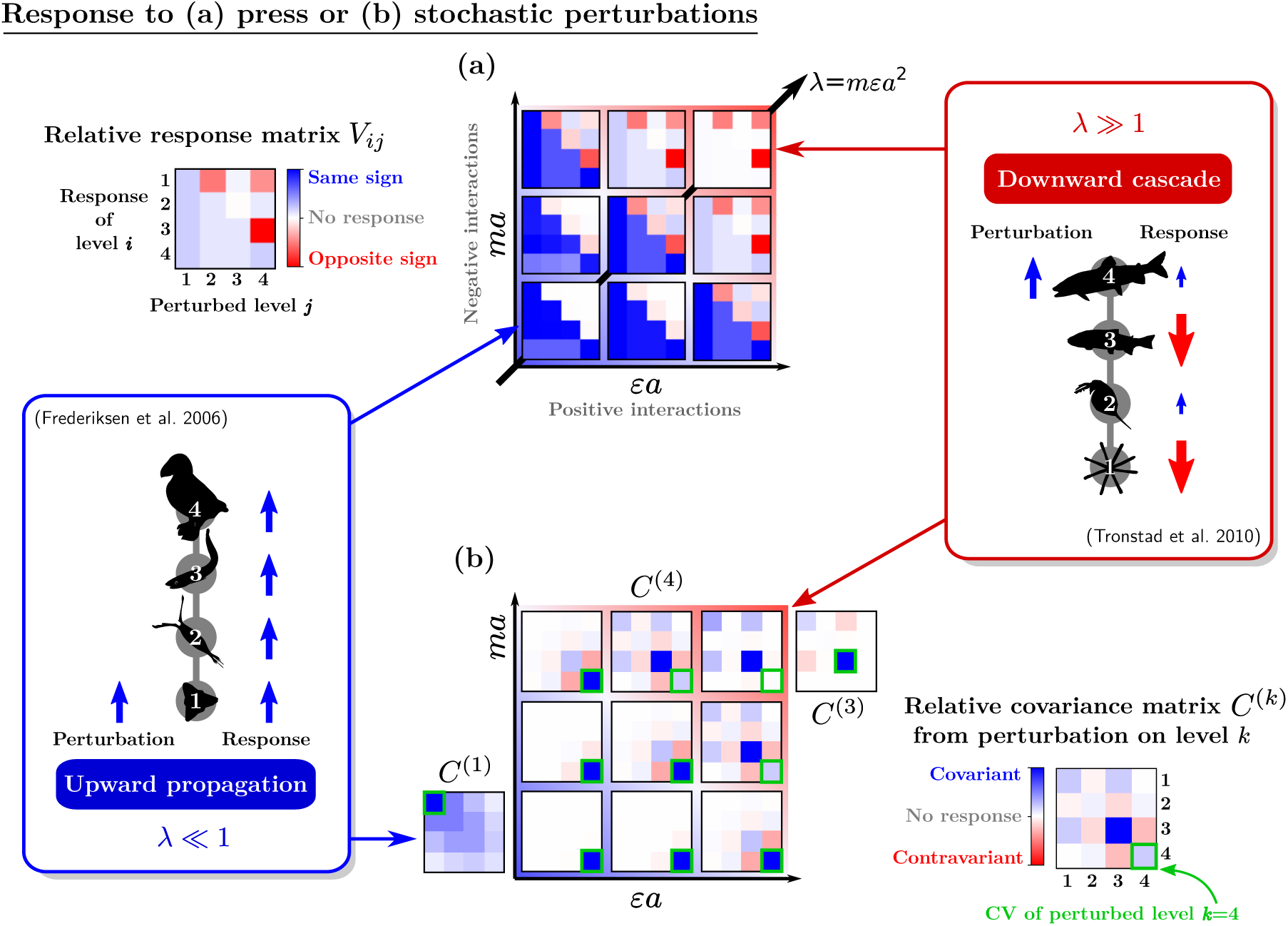
Map of responses to perturbations (Box 3). Some food chains display bottom-up patterns where variations in abundance are correlated across all trophic levels [52]. Other chains display top-down trophic cascades, with anticorrelations between adjacent levels [53]. These correlations can be measured in directional trends, resulting from press perturbations such as nutrient enrichment, or in undirected population fluctuations. **(a)** Long-term response to a press perturbation. The matrix *V*_*ij*_ measures the relative biomass change Δ*B*_*i*_/*B*_*i*_ in response to a relative change of growth rate Δ*g*_*j*_/*B*_*j*_. **(b)** Covariance of fluctuating time series. Applying a stochastic perturbation on one level *k* at a time gives a covariance matrix 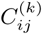, different for each perturbed level *k*. The diagonal element 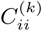 is the squared coefficient of variation (CV) of level *i*. The CV of the perturbed level is highlighted for comparison. In **(a)** and **(b)** colors (and arrows on the side diagrams) represent the sign and strength of the response, rescaled here by the largest coefficient of each matrix. We see that the main diagonal axis of the map, *λ* = *ma × εa* (the feedback of a level on itself through its predators), determines the nature of the response. Matrices *V* and *C*^(*k*)^ are both invariant along the other diagonal *κ* = *εa*/*ma*, which controls the chain’s top-heaviness in Fig. 3. This invariance does not hold for absolute stability metrics, see Fig. S1. In the bottom-up region (*λ* ≪ 1), the perturbed species is the one that responds most strongly, and perturbations only propagate upward, as illustrated by *C*^(1)^ and *C*^(4)^ where we affect either the basal or top level. In the top-down region (*λ* ≫ 1), a trophic cascade pattern (anticorrelated levels) is seen in *V* and *C*^(*k*)^. We also find an alternating pattern in the CV, with the top level being least variable and its prey being most variable, no matter which level is perturbed [54].

**FIG. 5.**
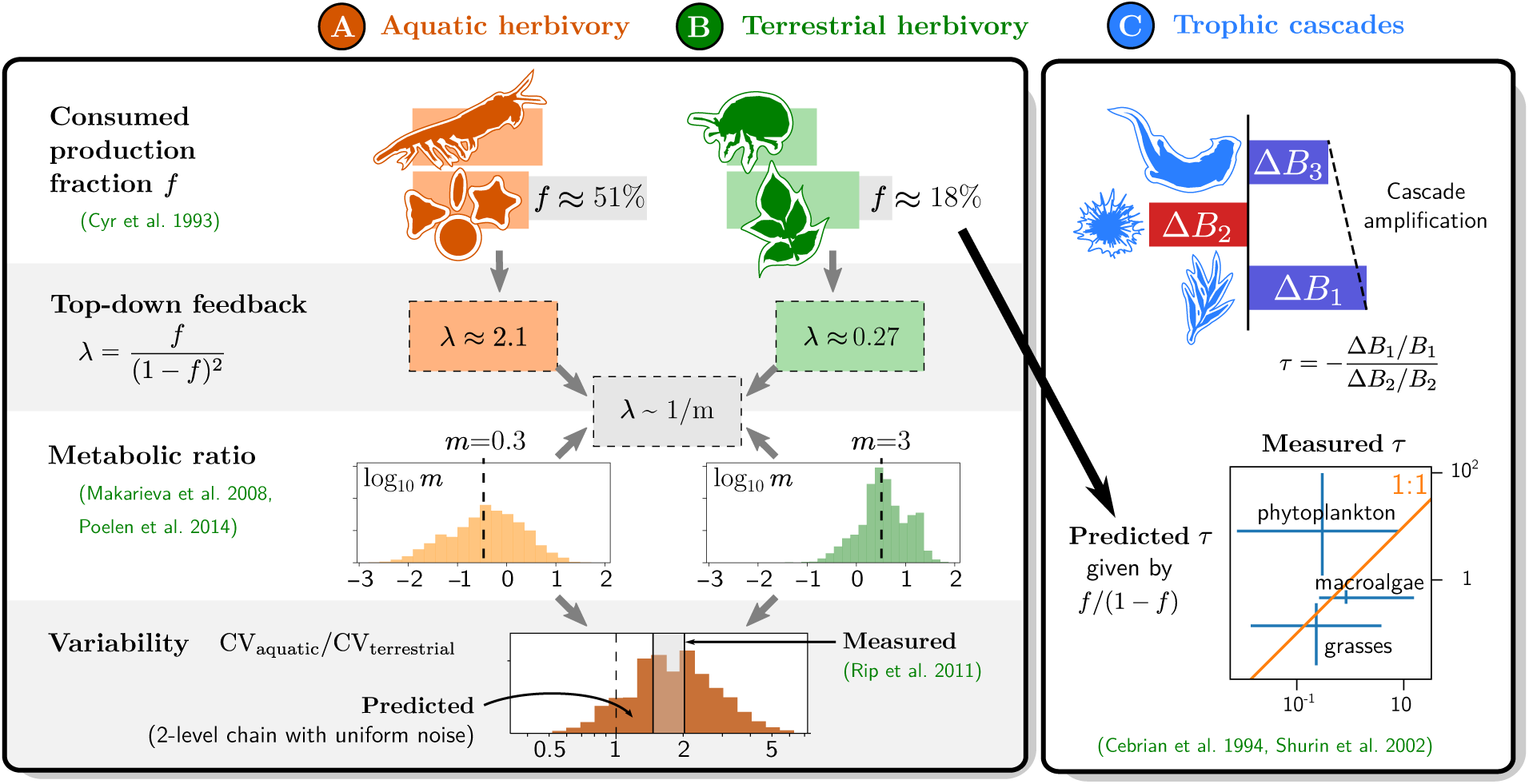
Empirical functioning and stability patterns, and their relationships predicted by the synthetic model. (A) Aquatic herbivory accounts on average for *f* = 51% primary production removal across various ecosystems, while (B) terrestrial herbivory only removes *f* = 18% of production [57]. They also differ by their median metabolic ratio *m*, measured from compiled data in Fig. S3 [46, 58]. The corresponding values of *λ* are compatible with the scaling *λ ∼* 1/*m* predicted by resource-driven consumption (Box 2). We then study stability properties (Box 3) using a two-level chain model with uniform noise. Parameterizing this model with *m* for each pair of species in the dataset, we obtain a distribution of CV for each ecosystem. We find that aquatic herbivores are less stable (higher CV) than terrestrial ones, in agreement with empirical data [59]. (C) Empirical measurements of the trophic cascade amplification factor *τ* in tritrophic chains [27], compared to predictions from data on removed production *f* for the basal level [42]. Measurements are grouped by the nature of the primary producer (bars are 25th to 75th percentiles), with phytoplankton-based chains exhibiting weaker cascades than expected from their large consumed fraction *f*. Despite considerable variance, we find indicative evidence of consistency between these various patterns.

We finally illustrate how this approach can lead to new insights into empirical data and quantitative tests of food chain theory. We discuss how these results may extend to more complex food webs and functional responses.

## A SYNTHETIC MODEL

### The energetic and dynamical paradigms

The energetic and dynamical paradigms described above appear to be at odds (Fig. 1), yet they can be understood as two different limits in a unified framework. We will demonstrate this with the minimal food chain model presented in Box 1.

We first establish a list of key ecological parameters, used throughout this study, which are common to both approaches. The environment determines the energy influx into the basal level, *g* [31]. Two important physiological parameters are the ratio *m* of predator and prey metabolic rates [30] and the efficiency *ε* with which consumed biomass is converted into growth [32, 33]. Finally, we must account for two types of ecological interactions: trophic interactions between levels, and self-regulation (e.g. direct competition) within a level [34, 35], which we denote by *α* and *D*, respectively. The ratio *a* = *α*/*D* plays a central role in our results, as it captures the relative strength of trophic and non-trophic feedbacks.

#### Box 1: Synthetic food chain model and key parameters

We propose the simplest model that can synthesize the predictions of both energetic and dynamical paradigms (Fig. 1 and 2).

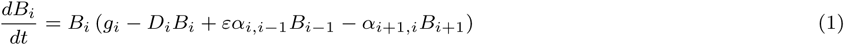

where *B*_*i*_ is the biomass of trophic level *i*, demographic processes are represented by intrinsic biomass growth or loss *g*_*i*_ and self-regulation *D*_*i*_, *ε* is the biomass conversion efficiency, and *α*_*ij*_ the strength of trophic interactions. All the rates appearing in this equation may depend on the species’ metabolic rate *m*_*i*_. For simplicity, we assume a linear scaling:

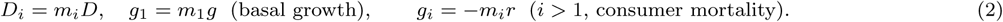

Here, *g* represents the basal energy influx, including nutrient supply and autotroph efficiency, while *r* represents biomass loss and mortality due to metabolic costs. Note that the carrying capacity *g*_1_/*D*_1_ of the autotroph level sets the scale for the total biomass in the chain. To allow direct comparison between density-dependent and independent terms, we can choose biomass units such that *g*_1_ = *D*_1_. The model further simplifies if the predator-prey metabolic ratio and the ratio of interaction strength to self-regulation

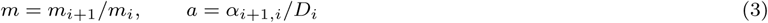

are both constant throughout the chain. The equilibrium condition for surviving consumers (*i* > 1), which we derive and solve in Appendix S1 in Supporting Information, is then

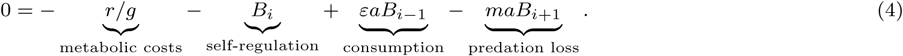

Its solution is the equilibrium biomass distribution, which depends on three synthetic parameters:

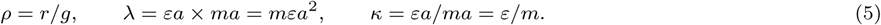

The product *λ* denotes the strength of top-down control in the chain. It represents the feedack of a trophic level on itself through its predators: how much one unit of prey biomass increases predator biomass at equilibrium (*εa*), times how much this additional predation suppresses the prey (*ma*), normalized by the prey’s self-regulation. The ratio *κ* represents how much biomass is gained by consumers per unit biomass lost by resources, and we see in Fig. 3 that large *κ* allows top-heavy distributions. Finally, *ρ* captures the balance between metabolic losses *r* and basal energy influx *g*. Losses can limit growth or even cause extinctions, but if *g* is comparatively large (*ρ ≈* 0), their effect becomes negligible, and equilibrium patterns are instead determined by interactions alone.

The two paradigms share a basic account of biomass creation, loss and transfer between levels [16]. Given the biomass *B*_*i*_ and production *P*_*i*_ (biomass created per unit time) of trophic level *i*, we can write the dynamical and static equations

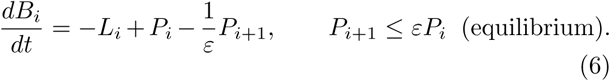

where *ε* is the conversion efficiency defined above, and *L*_*i*_ represents all non-trophic biomass losses. The static inequality becomes an equality when *L*_*i*_ = 0. Since *ε* < 1, any food chain at equilibrium must display smaller energy flux (production) at higher levels.

The energetic paradigm emphasizes static relationships such as the equilibrium condition in (6). To predict biomass distributions, various studies [36–38] have proposed heuristic equations for energy *stocks* rather than *fluxes*:

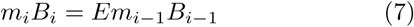

where *m*_*i*_ is the metabolic rate of trophic level *i*, and *m*_*i*_*B*_*i*_ is interpreted as its energy content. Some studies identify fluxes and stocks, *P*_*i*_ = *m*_*i*_*B*_*i*_ [39], and thus *E* = *ε*. In general, this identification does not hold and *E* is not an efficiency, but a phenomenological factor [36]. Our model yields the relationship *E* = *εma* (Appendix S1) and we note that, contrary to fluxes, no physical principle prevents the accumulation of larger energy stocks at higher levels, *E* > 1.

By contrast, the dynamical paradigm emphasizes the nonequilibrium patterns arising in (6) from feedbacks between predator and prey. Behavior and physiology are encapsulated in a functional response [40] specifying how consumer production, *P*_*i*_, depends on consumer biomass, *B*_*i*_, and resource biomass, *B*_*i*−1_. To facilitate comparisons, we follow a classic model with metabolic scaling [30] and adopt a simpler Lotka-Volterra (Type 1) functional response, giving

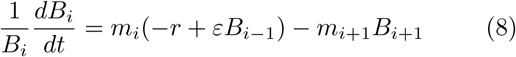

where *r* represents metabolic costs. This corresponds to setting production *P*_*i*_ = *εm*_*i*_*B*_*i*_*B*_*i*−1_ and losses *L*_*i*_ = *rm*_*i*_*B*_*i*_ in (6). More complex models are considered in the Discussion.

Our central observation is that the energetic formula (7) cannot arise as an equilibrium of the dynamical equation (8), but that it could with the addition of self-regulation. We construct our synthetic model (Box 1) as a dynamical equation with self-regulation, and show in Fig. 2 that it admits both (7) and (8) as special cases (we show in Appendix S1 that this holds for other ways of modelling self-regulation, e.g. predator interference). This allows a quantitative comparison between energetic and dynamical models.

### Connecting the paradigms

We show in Fig. 2 how the synthetic model developped in Box 1 connects with both paradigms.

The first discrepancy between the two paradigms lies in the role of self-regulation, which is central, if implicit, in the energetic argument, yet often absent from dynamical models. We will see that these two choices correspond to opposite extremes in the value of the synthetic parameter *λ* = *mεa*^2^, which encapsulates the balance between self-regulation and predation losses (Box 1).

This discrepancy may reflect the difference between a microscopic viewpoint, where a predator species may dramatically deplete its prey species in a small locality, and a macroscopic viewpoint, where we generally find stable coexistence of trophic levels [41]. Yet, even locally, empirical evidence suggests the underestimated role of self-regulation, notably in the form of predator interference [35].

#### Box 2: Metabolic scaling of interactions

In the model defined in Box 1, interactions *α*_*i*+1,*i*_ can depend on both consumer metabolism *m*_*i*+1_ and resource metabolism *m*_*i*_. The dynamical paradigm assumes that attack rate is proportional to consumer metabolism, hence *α*_*i*+1,*i*_ ∼ *m*_*i*+1_. In the energetic paradigm, energy transfer is proportional to the resource’s energy content (see Fig. 2). This implies that interactions scale with resource metabolism, *α*_*i*+1,*i*_ ∼ *m*_*i*_. These two choices can be summarized in a generalized equation, with a parameter *ν* = *−*1 (resource-driven) or *ν* = 0 (consumer-driven):

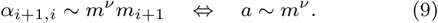

Note that *λ* = *mεa*^2^ ∼ *m*^1+2*ν*^. In the dynamical paradigm (*ν* = 0), top-down control increases with metabolic ratio, *λ ∼ m*. In the energetic paradigm (*ν* = *−*1), top-down control decreases with metabolic ratio, *λ* ∼ 1/*m*. We discuss the empirical evidence for each scaling in the main text (see Fig. 5).

The second difference is that energetic models generally assume negligible energy loss through metabolic costs, *ρ* ≪ *r*/*g* 1 (Box 1). In that limit, basal energy influx *g* plays no role in the dynamics, only acting as a constant factor in the equilibrium biomass of all levels. The dynamical paradigm, however, assumes significant losses and thus assigns a major role to basal energy influx *g*, which can determine whether higher trophic levels go to extinction. This can notably lead to the paradox of enrichment [26], where an increase in nutrient supply may cause a loss of stability.

The third difference lies in the metabolic scaling of interactions (see Box 2 and Fig. 2). Dynamical models (8) again adopt the consumer’s perspective: a predator with a faster metabolism is expected to have a higher attack rate, hence the consumption rate *α*_*i*+1,*i*_ scales with the *consumer*’s metabolism *m*_*i*+1_ [30]. On the other hand, energetic models assume that consumption – or energy transfer, the right-hand term in (7) – scales with the metabolic rate of the *resource*.

As a result of these conflicting assumptions, we expect the two paradigms to emphasize either bottom-up or top-down effects, but also make different predictions about the effects of nutrient enrichment and metabolic scaling.

### Bottom-up and top-down control

We now explain how the assumptions of the energetic and dynamical paradigms lead to a prevalence of bottom-up or top-down patterns, respectively, in the food chain. We also point out that the two paradigms have conflicting notions of bottom-up control.

Consider a two-level chain as an illustration. In the absence of any self-regulation, setting *D*_*i*_ = 0 (and metabolic scaling *α*_*i,i*−1_ = *m*_*i*_*α*) in (1), the chain reaches the equilibrium

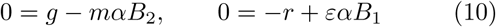

where we see that each level controls the other. Resource biomass *B*_1_ is fixed by consumer mortality *r*, corresponding to top-down control (if *r* is small, the resource is almost driven to exhaustion). On the other hand, consumer biomass *B*_2_ is fixed by the basal energy influx *g*. While consumers are limited by a constraint from below (and this is sometimes called a bottom-up effect), this is a consequence of their own control of the resource, as they grow until they divert all of the resource’s production to maintain their biomass.

This situation persists if at least one trophic level lacks self-regulation: whether consumers or resources, that level will grow until it is limited by the antagonistic interaction, i.e. until consumers remove all extra resource growth and cannot grow further themselves.

It is only with self-regulation *D*_*i*_ > 0 at *both* levels that the two populations can stabilize before resource production is entirely consumed. Self-regulation thus gives rise to the classic notion of *donor* control, where a trophic level’s biomass is fixed by its prey’s, but does not increase enough to exert a negative impact on that prey. This low-impact coexistence of predator with prey cannot happen if even a single level lacks self-regulation.

An important difference between these two settings is the relationship between biomass *B*_*i*_ and production *P*_*i*_ or productivity *p*_*i*_ = *P*_*i*_/*B*_*i*_, widely studied empirically [22, 42] and illustrated in Fig. 1.

*Donor* control is characterized by the fact that a level’s production determines its own equilibrium biomass *B*_*i*_. In our model (Box 1), per-capita losses from self-regulation are proportional to *B*_*i*_, and when they equilibriate with per-capita growth *p*_*i*_, we obtain *B*_*i*_ ∼ *p*_*i*_.

On the other hand, *antagonistic* control is characterized by the fact that each level’s growth or losses determines the other’s biomass. All prey production goes to predators, hence *B*_*i*_ ∼ *p*_*i*−1_. Counter-intuitively, there is no correlation between a trophic level’s biomass and its own productivity or mortality, as we see in (10).

Following usage in empirical studies, “bottom-up” will hereafter be used in the donor sense, rather than in the antagonistic sense in which top-down and bottom-up control co-occur as two sides of the same interaction.

## FUNCTIONING PATTERNS

We now use our model to clarify theoretical relationships between food web patterns, illustrating them for a four-level food chain in the limit of low mortality or high basal energy influx *ρ* ≪ 1. This limit ensures that trophic levels cannot go extinct, and that our description is robust to changes in food chain length or primary production. We show in Appendix S1 how our results can be extended to arbitrary mortality rates.

### Biomass and production

One of the most central predictions of food chain theory is the distribution of energy among trophic levels, in the form of either biomass or growth. Since Elton [1], energetic arguments have been used to predict pyramidal or hierarchical patterns (Fig. 3), but few studies have investigated when this structure can emerge dynamically [28, 44, 45].

We see in Fig. 3a that biomass patterns vary along two dimensions: *ma* and *εa*, the strength of negative and positive interactions in (4) (Box 1). To better understand how biomass is distributed, we must combine these two quantities, taking either their product or their ratio,

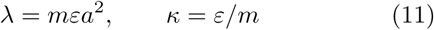

i.e. the diagonals represented on Fig. 3b.

As shown in Box 1, the product *λ* denotes the strength of predation feedback, i.e. how much a trophic level limits itself through its predators. The ratio *κ* indicates whether biomass accumulates toward the top or the bottom of the chain. Notice in (4) that if we can neglect predation losses, then 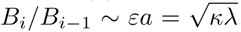, whereas if we can neglect self-regulation, then *B*_*i*+1_/*B*_*i*−1_ ∼ *ε*/*m* = *κ*. In both limits, larger *κ* leads to more top-heavy chains.

Between these two limits, a useful proxy for the importance of top-down control is

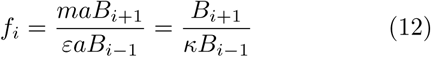

which can be interpreted as the fraction of production at level *i* lost to consumer *i* + 1.

If we assume a pyramidal structure, i.e. a constant biomass ratio between adjacent levels *B*_*i*+1_/*B*_*i*_, then *f*_*i*_ must also be constant and we get the scaling

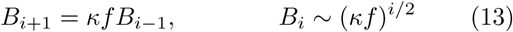

This is indeed a possible solution of the equilibrium equation (4) with *f* given by

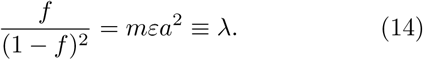

Thus, *λ* and *κ* together define the trend of the biomass pyramid (13) toward bottom-heaviness (*κf* < 1) or top-heaviness (*κf* > 1), illustrated in Fig. 3.

Yet we also expect this pyramidal structure to disappear for strong top-down control, i.e. low self-regulation, as in (10). For *λ* ≳ 1, a cascade pattern emerges. Starting from the top level, the chain alternates between high and low-biomass levels (Fig. 3a), controlling each other through their antagonistic interaction. High-biomass levels remove a large fraction *f* of their prey’s production, and they still follow the scaling (13), which we call the global slope of the cascade. With increasing *λ*, low-biomass levels converge toward the same scaling, with a smaller prefactor.

We can show (see Appendix S1) that the biomass ratio between high- and low-biomass levels is 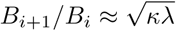. Since *f ≤* 1, for large *λ* this ratio can become much larger than the global slope 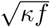. Thus, if one can only observe two adjacent trophic levels, their biomass ratio may give the illusion of a steep inverted pyramid, as in the empirical example in Fig. 3c discussed below.

### Biomass-size scaling

The distribution of energy among trophic levels is more readily observed when these levels are clearly distinct, e.g. in simple plant-herbivore-carnivore chains. In many systems, however, trophic height is not so easily assessed. Instead, the most immediate property of an organism is often its size, especially in aquatic food webs. The study of these ecosystems thus relies on the distribution of abundance or total biomass per size class [18, 21].

Assuming a fixed relationship between size and trophic height, the pyramidal slope (13) can be translated into a continuous distribution as a function of body mass (or size) *W*_*i*_. Since the metabolic rate *m*_*i*_ is also less accessible than body size, many studies posit an allometric scaling of metabolism with size, 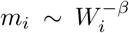 [30] with some exponent *β* measured empirically.

Let us define the predator-prey body mass ratio *M*, which is assumed to be constant throughout the chain:

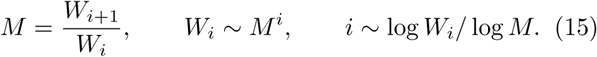

We thus have an expression for the trophic level *i* for size class *W*_*i*_. By substituting this into the exponent in (13) and using the allometric scaling *m ∼ M^−β^*, we find the biomass-size scaling for a continuous variable *W*

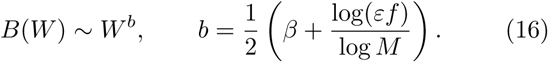

We can deduce the number of individuals per size class, *B*(*W*)/*W*, also known as size spectrum [18, 19]. Previous studies [36] have used the energetic argument (7) and obtained a similar prediction, but with a different exponent: *b* = *β* + log *E*/log *M*. Our formula extends it with two terms coming from the predation feedback: the fraction of production removal *f*, and the prefactor 1/2. The two formulas coincide if *f* 1 (*f* ≈ *λ*) as shown in Appendix S1.

From metabolic data, it is generally estimated that *β* ∈ 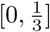 [46–48]. While top-heavy pyramidal distributions are allowed by (16), they are expected to have at most exponent 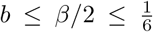 (given *ε, f* ≤ 1 and assuming that consumers are larger, *M* > 1). For instance, in an oceanic food chain, large mammal biomass could at most exceed unicell biomass by two orders of magnitude. In fact, it has been proposed that *B*(*W*) is approximately flat, *b* ≈ 0, over many orders of magnitude in marine data [21].

A steeper slope in the scaling may only be seen over a limited size range, between a low-biomass level and a high-biomass level in a cascade pattern, *B*_*i*+1_/*B*_*i*_ ≈ 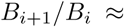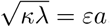, giving

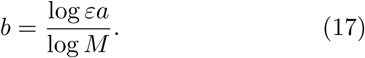

We now show that we can use the biomass-size exponent to probe the structure of an empirical food chain.

### The “paradox” of positive biomass-size scaling

Fish in kelp forests exhibit a top-heavy biomass distribution per size class, as shown in Fig. 3c. A recent study [43] described this pattern as a paradox, because its slope appears steeper than allowed by energetics: the authors find *B*(*W*) ∼ *W*^*b*^ with exponent *b* ≈ 0.4, suggesting an impossible efficiency *ε* > 1 in formula (16).

To solve this paradox, we first notice that the exponent *b* is measured over a limited size range, spanning less than two trophic levels. Indeed, the study uses stable isotope analysis to estimate the mass ratio between trophic levels, but the observed spectrum covers only body masses from 32 to 2048g (a ratio of 64 between the largest and the smallest fish). Thus, it may be misleading to analyze this spectrum as a multi-level biomass pyramid described by formula (16).

Instead, we suggest a top-down cascade pattern (Fig. 3c), with the largest fish in the sample belonging to a high-biomass level. In that case, equation (17) yields *εa* ≈ 28, suggesting trophic interactions far stronger than self-regulation. We thus expect strong trophic cascades, which could be tested by predator removal. It is also important to note that the assumption of constant mass ratio *M*, used by [43] to convert between size classes and trophic levels, may be problematic as *M* is very widely distributed in different interactions (Fig. S4).

#### Box 3: Stability properties

We study the quantitative stability properties of the model in Box 1, i.e. its response to various types of perturbations. The dynamical equation (1) becomes

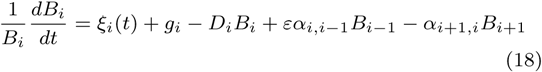

where *ξ*_*i*_(*t*) represents the perturbation.

We focus on two types of long-term perturbations. In the first case, we compute the effect of a constant press perturbation *ξ*_*i*_(*t*) = *ξ* on one trophic level at a time. This can be interpreted as a permanent change Δ*g*_*i*_ of its growth or mortality term, up to complete removal of the level. We predict the resulting change in abundance Δ*N*_*j*_ at each level.

In the second case, we add demographic stochasticity to one level at a time, 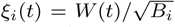 with *W* (*t*) a white noise term (see Appendix S1). For each choice of the perturbed level, we then compute the covariance matrix of the biomass fluctuations of all levels. On the diagonal of this matrix, we find the variance of each level. The inverse of its trace, *invariability*, has been widely used as an empirical stability measure [49]. These two properties are connected to other stability properties, such as structural stability and return time to equilibrium [50, 51].

In Fig. 4, we show results for *relative* stability metrics, rescaled by equilibrium abundances 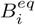:

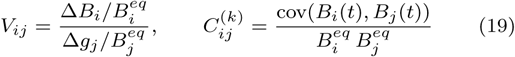

with *V*_*ij*_ the relative response of level *i* to a press on *j*, and *C*^(*k*)^ the relative covariance matrix between all levels created by perturbing level *k* only. The diagonal element 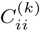 is the coefficient of variation (CV) of level *i*. This rescaling reveals clear pyramidal or cascade patterns that are essentially determined by a single parameter, *λ*. When using absolute metrics (Fig. S1), these patterns are skewed by the biomass distribution and less apparent to the eye.

## STABILITY PATTERNS

### Trophic cascades and response to a press

Trophic cascades are studied by comparing systems where the top predator is absent or present, e.g. [55], or modifying its abundance by a smaller amount.

More generally, we consider a long-term decrease or increase in any trophic level’s biomass, due for instance to harvesting or nutrient enrichment. It can be modelled as a press perturbation (Box 3), equivalent to a change Δ*g*_*i*_ of the growth or mortality rate, *g*_*i*_, of one trophic level in (1). We then study the matrix of the relative responses Δ*B*_*j*_/*B*_*j*_ to a press Δ*g*_*i*_/*B*_*i*_, shown in Fig. 4a.

Upward propagation of perturbations, whereby a decrease (increase) in the abundance of a trophic level leads to a decrease (increase) at all higher levels, appears as a blue lower triangle in the press response matrix in Fig. 4a. This dominates the community response when *λ* < 1. Downward cascades are characterized by alternating negative and positive responses of the levels below the perturbed level, which appear in alternating red and blue in the upper triangle of the response matrix. They dominate when *λ* > 1. This coincides with the qualitative shift in the shape of the biomass distribution (Fig. 3), illustrating the tight connection between patterns in biomass and patterns in stability, see e.g. [54].

We show here these patterns when the food chain is not limited by energy loss (small *ρ* = *r*/*g*, see Box 1), and cascade strength only depends on *λ*. In energy-limited food chains (large *ρ*), the picture is more complex. In Appendix S1, we compare a trophic level’s biomass before and after predator removal, and show that, depending on all dynamical parameters, both increase with basal energy flux *g* but either may increase faster. There is no simple prediction for how *g* (e.g. nutrient enrichment) affects trophic cascades: when it is large, cascade strength is independent from it, and when it is small, the dependence may go in either direction.

### Variability under stochastic perturbations

Bottom-up and top-down effects can also be measured as correlations in empirical time series [52, 53]. Correlative measures capture directional trends, such as the press response discussed above, as well as undirected fluctuations. These fluctuations also contain information about the food chain structure.

In the model, we can impose stochastic noise on one level and measure the variances and covariances of all levels’ time series (Fig. 4b). We rescale them by equilibrium biomasses to obtain a relative covariance matrix (Box 3). The diagonal elements of this matrix are the squares of the coefficient of variation (hereafter CV) for each trophic level.

For low *λ* ≪ 1, the trophic level with the highest CV (shown on the diagonal of the relative covariance matrix) is the one that is directly perturbed, and this perturbation propagates only to higher levels, which covary positively.

For intermediate and large *λ* ≳ 1, we see a distinct checkerboard pattern (covariance cascade) for all levels below the perturbed one, reflecting the tendency of prey abundaces to fluctuate in opposition to their predators.

We also note that, deep in the top-down regime *λ* ≫ 1, the level with the lowest CV is always the top predator, while its prey has the highest CV, and lower levels alternate between the two, similarly to theoretical patterns in production and biomass [3, 4, 54]. Perturbing intermediate levels (see *C*^(3)^ in Fig. 4b) gives rise to a “two-step” covariance cascade: species two levels apart exhibit anti-correlated fluctuations. This can be interpreted as a cascade between predator-prey pairs, rather than between individual levels, due to the prey being tightly controlled by its predator.

In a dynamical model with a nonlinear functional response, these anticorrelated fluctuations around equilibrium could announce the transition to more complex dynamical regimes such as predator-prey cycles, or chaos, as we note in Discussion.

## EMPIRICAL APPLICATIONS

We now illustrate how the synthetic model can be used to interpret and connect diverse empirical phenomena.

Theoretical patterns, from pyramids to cascades, have been identified above under the assumption that the parameters *λ* and *κ* are constant throughout the chain, due to equations (2) and (3). These idealized patterns provide meaningful intutions, but they can be distorted by large parameter variations between trophic levels [28, 44].

In empirical settings, we do not expect all trophic levels and interactions to follow this simple parameterization. Different ecological processes need not follow the same scaling with metabolic rates, and most food chains involve organisms belonging to distinct taxonomic classes, with vastly different physiologies.

We compiled data on thousands of predator-prey pairs (Fig. S3 to S5) and found that the metabolic ratio *m* varies over four orders of magnitude, *m ∈* [10^−2^, 10^2^]. While the body size ratio *M* displays a clear skew toward predators being larger than their prey, there is no such skew in metabolic ratio: faster and slower consumers are equally common in the dataset, with significant variation between and within taxonomic classes.

Biomass conversion efficiency *ε* has a smaller range of variation, often estimated around 10 *−* 20% [32, 33, 38, 56], but it may still differ between trophic levels [30], and we note in Discussion the issues with quantifying interaction strength *α* and self-regulation *D*.

In the following, we thus allow different parameters for each trophic level, and show how the model can be parameterized using empirical evidence to produce quantitative predictions.

### Differences between aquatic and terrestrial herbivory

Food web ecology has long focused on explaining dissimilarities between aquatic and terrestrial food webs [7]. One striking difference is that aquatic herbivores consume on average *f* = 51% of primary production across many ecosystems, while terrestrial herbivory only removes *f* = 18% of plant production [57]. This discrepancy has been explained by terrestrial autotrophs having slower turnover than their consumers, while aquatic autotrophs, especially phytoplankton, have much faster turnover [7]. In some systems, this allows aquatic herbivores to consume close to 100% of phytoplankton biomass and production each day [42].

The consistency of this explanation can be tested quantitatively in our model, as we show in Fig. 5. Using measurements on *f*, we can compute *λ* = *f*/(1 − *f*)^2^. Then, using the median metabolic ratio *m* for either aquatic or terrestrial herbivory in the metabolic data (Fig. S3 to S5), we check that *λ* ∼ 1/*m*, in agreement with the scenario of resource-driven consumption (Box 2, energetic paradigm).

Furthermore, aquatic herbivore populations have been shown to be less stable (higher CV, defined in Box 3) than terrestrial herbivores [59]. Using the scaling *λ* ∼ 1/*m* and the data for herbivore-plant metabolic ratios *m*, we have computed herbivore CV for each pair in the simplest theoretical setting (a two-level chain with equal noise on both levels). Doing all possible pair comparisons between aquatic herbivores and terrestrial ones, we find that the former tend to have higher CV. The aquatic-terrestrial CV ratio is widely distributed but centered around the empirical range [1.5,2.1] found in [59].

Hence, functioning and stability differences between aquatic and terrestrial food chains could both have a physiological origin in metabolic rates, consistent with the metabolic scaling assumed in the energetic paradigm, but accounting for predation mortality.

### Cascade strength in tritrophic chains

Trophic cascade strength is commonly measured as the logarithmic change (log ratio) in plant biomass Δ log *B*_1_ in response to predator manipulation [60]. It is related to the relative response shown in Fig. 4a by Δ*B*_1_/*B*_1_ = exp(Δ log *B*_1_) − 1.

In a meta-analysis, the plant biomass log ratio was found to exhibit strong positive correlation to the carnivore’s metabolic rate, *m*_3_, and negative correlation to herbivore’s, *m*_2_ [10]. This correlation can be predicted in a three-level chain (see Appendix S1) under two conditions: first, the carnivore must have significant self-regulation *D*_3_; second, its attack rate must scale with its own metabolic rate, *α*_32_ *∼ m*_3_, in agreement with consumer-driven consumption (Box 2, dynamical paradigm) and in contrast with what was found above for herbivory.

Another salient property of cascades is whether the effect attenuates or intensifies down the chain, e.g. whether plants are less or more affected than herbivores by carnivore removal [6, 61]. This factor has been measured empirically, and explained by different biological mechanisms (from plant defenses to external subsidies) in a variety of systems. In the three-level chain, we find

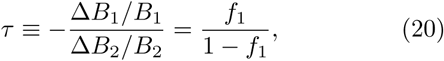

indicating that, without the need to invoke additional mechanisms, attenuation happens if the fraction of consumed primary production is less than half, *f*_1_ < 50%, while intensification happens otherwise.

From data on herbivory [57], we estimate an amplification factor *τ* ≈ 1 in aquatic foodwebs, meaning that cascade effects tend to propagate without attenuation. By contrast, we expect fast-decaying cascades with *τ* ≈ 0.2 in terrestrial systems. In Fig. 5, we use additional data on *τ* from [27] and *f* from [42] to show that these predictions are plausible, although with large variance. Combined with the previous section, this is indicative evidence that metabolic rates may cause the main trend in trophic cascade attenuation.

### The stabilizing role of metabolic scaling

Previous studies have emphasized a possible stabilizing role of allometric scaling [30]. Since predators are larger than their prey, allometric scaling suggests that they should have slower metabolism. With a median size ratio *M* ≈ 80 [62], 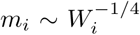 gives a metabolic ratio *m* ≈ 0.3. These studies have shown that this favors stability and coexistence in dynamical simulations of complex food webs.

Our food chain model can reach a similar or opposite conclusion, with a simple explanation. Consumer-driven consumption (*α*_*i*+1,*i*_ ∼ *m*_*i*+1_) gives *λ* ∼ *m*, as noted in Box 2. Therefore, slower predators with lower *m* induce weaker predation feedback *λ*, which leads to greater stability (Fig. 4), in qualitative agreement with the simulation studies. This scaling is supported by observations on cascade strength in the previous section.

On the other hand, for resource-driven consumption (*α*_*i*+1,*i*_ ∼ *m*_*i*_), we reach the opposite conclusion: *λ* ∼ 1/*m* and slower consumers *destabilize* the system. This is supported by the fact that aquatic herbivores have both lower metabolic ratio *m* and higher variability than terrestrial ones (Fig. 5).

These two conflictual lines of evidence challenge any simple and universal stability argument for metabolic scaling [30]. We note that, despite predators being generally larger than their prey, there is no global tendency for them to have a slower metabolism, as we show in data compiled from multiple studies (Fig. S3 to S5). Therefore, it may be that different ecological settings favor either slower or faster consumer metabolism.

## DISCUSSION

Our understanding of many ecological phenomena relies on intuitions developed from simple food chains. Despite its fundamental role, this body of work has become increasingly fragmented. First, functioning and stability patterns, such as biomass pyramids and trophic cascades, have become disjoint topics. The former are often analyzed with energetic arguments, and the latter with dynamical models (Fig. 1). But these approaches are based on conflicting assumptions, as we clarified by embedding them in the same formalism (Fig. 2).

Second, food chain behaviors arise from the interaction of multiple ecological parameters: basal energy influx *g*, predator-prey metabolic ratio *m*, biomass conversion efficiency *ε* or interaction strength *a*. Many empirical and theoretical studies have investigated a single axis at a time, occasionally reaching contradictory conclusions. We have shown here that the action of these key parameters can only be understood in conjunction.

We have summarized the main food chain patterns, and the parameters that control them, in a concise map with two main axes. Its first axis is given by the synthetic parameter *λ*, which denotes the strength of top-down control in the chain, as it quantifies the feedack of a trophic level on itself through its predators (Box 1).

The bottom-up regime (*λ* < 1) is characterized by pyramidal patterns, regular or inverted, in various functioning and stability properties (Fig. 3 and 4). In the top-down regime (*λ* > 1), alternating cascade patterns are found instead.

The second synthetic parameter, *κ*, controls the overall top-heaviness of the biomass distribution, but has no effect on essential stability properties (Fig. 4). Counter-intuitively, top-heavy distributions do not imply strong top-down control: species with more efficient assimilation and slower turnover can acquire and store larger amounts of energy, without necessarily exerting stronger pressure on lower levels.

A third parameter, *ρ*, modulates these patterns without creating qualitatively new regimes. It represents the fraction of energy lost to mortality and metabolic costs, and is important only if these losses significantly reduce a trophic level’s biomass. When *ρ* ≪ 1 (low metabolic costs *r* or high basal energy influx *g*), its precise value becomes irrelevant to food chain dynamics.

By combining the ingredients of the energetic and dynamical paradigms (Fig. 2), the synthetic approach explains their discrepancies, recovers their main results, and extends them beyond their traditional scope.

These results provide an intuitive basis for understanding relationships observed in data or complex simulations, and advancing our understanding of a number of standing empirical paradoxes. We have shown that they could provide new quantitative predictions on relationships between consumption, metabolism and trophic cascade strength.

### Self-regulation

We have shown that self-regulation is at the heart of the main disagreement between the energetic and dynamical approaches (Fig. 2). By preventing consumers from growing until they divert all of their resource’s production, self-regulation stabilizes the dynamics [63], attenuates top-down cascades, and gives rise to bottom-up pyramidal patterns, where each level’s dynamics depends on its energy influx only.

Few studies reliably quantify trophic interaction strength *α* [34], and fewer provide estimates of self-regulation *D* [35]. But we have shown in Fig. 5 that indirect estimates of their ratio *a* = *α*/*D* can be obtained from various empirical patterns, allowing us to evaluate the relative importance of self-regulation.

Many food web models, e.g. [30], do not admit explicit self-regulation in the form of density-dependent mortality *D*_*i*_, except at the basal level. Nevertheless, they often contain predator interference [64] which is widely supported by empirical evidence [22, 35].

Interference plays a similar self-regulating role at equilibrium. For instance, this can be seen in the dynamics of a consumer with a Type II functional response and predator interference,

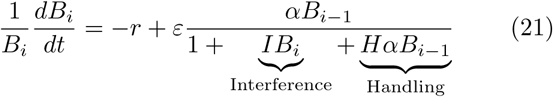

where *I* is the interference strength and *H* is the handling time. At equilibrium, we find

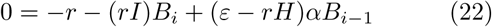

which is identical to the top predator’s equilibrium in our Lotka-Volterra model (1) with self-regulation *D* = *rI* and a reduced conversion efficiency *ε* − *rH*.

The limit of strong interference, where the denominator in (21) reduces to *IB*_*i*_, has been widely discussed as a *ratio-dependent* functional response [65]. We show in Appendix S1 that it also allows a pyramidal biomass distribution. We interpret ratio-dependence and density-dependent mortality as two examples in a wider range of self-regulation mechanisms, and expect that our qualitative results may extend to other forms of density dependence stabilizing the dynamics [22, 66].

### Disentangling effects of primary productivity

This synthetic approach sheds a new light on the effects of primary productivity on food chain behaviors [26, 59]. In Box 1, we defined primary productivity as *g*_1_ = *g m*_1_, where *g* is tied to nutrient supply and *m*_1_ to autotroph metabolism. There are thus two ways in which ecosystems can differ in their primary productivity.

Nutrient enrichment increases the basal energy influx *g*. All else being equal, this will increase the total biomass, but produce inconsistent effects on stability and trophic cascades. We show in Appendix S1 that these effects can be positive or negative depending on the other parameters, and they vanish at high values of *g*, when growth is not limited by metabolic losses.

On the other hand, an increased metabolic rate *m*_1_ can have a consistent effect on stability. The evidence in Fig. 5 suggests that when primary producers have faster metabolism than their consumers, as in aquatic systems, one finds stronger trophic cascades and more variability.

This provides a tentative explanation for conflicting empirical findings on the effects of primary productivity, especially in cross-ecosystem comparisons where *g*, *m*_1_ and other parameters may all vary [5, 10].

### Application to real food webs

Our idealized food chain model trades realism for ease of interpretation and mathematical treatment. Yet, a complex web structure is an ubiquitous feature of real trophic interactions, and nonlinear functional responses (how consumption depends on the density of predator, prey, and even third parties) can be added to capture physiological and behavioral characteristics [40].

We expect these two ingredients to have predictable effects in the region of bottom-up control, but far more complex ones in the region of top-down control.

When self-regulation greatly exceeds predation losses (*λ* ≪ 1), there are no complex dynamical feedbacks, as each species determines its consumers’ abundance but is almost unaffected by them. In that case, food chain models can easily be amended to account for additional fluxes, e.g. for omnivorous predators [15] or structured populations [38]. Different functional responses affect quantitative results, but may not lead to qualitatively different dynamical regimes.

For weak self-regulation (*λ* ≫ 1), however, food web dynamics may become excitable and lead to cycles or chaos, unless species satisfy complex conditions [67–69]. Even qualitative functioning and stability properties become sensitive to interactions motifs and functional responses, as described in a vast literature [9, 25, 70–72], and it is challenging to extend our near-equilibrium results to these nonequilibrium dynamics.

Nevertheless, coherent top-down patterns, such as trophic cascades, are well-attested in various ecosystems [5, 73]. This suggests that our simple approach remains valid in some limits: either for particular sets of species, or for averages over large communities. One might be able to unfold a complex web into its essential chain-like structure [74] to parameterize our model from more realistic descriptions.

### Conclusions and prospects

While static energy pyramids, dynamical fluctuations and trophic cascades have been studied with different approaches, they all arise from the same food chain structure. We have synthesized these basic predictions in a simple two-dimensional map, whose axes combine physiological and ecological parameters.

This map is divided in two regions, in which either pyramidal or cascade patterns can be found across a wide range of stability and functioning properties. This dichotomy reflects two intrinsically different dynamical regimes, one dominated by donor control and the other by antagonistic feedbacks. But rather than extremes on a continuum, these regimes have often been approached as alternative ways to understand and model food chains, each ingrained in a long tradition and associated with its own set of questions and methods.

We therefore emphasize the need for consistency between the results of these different approaches. Considerable empirical and theoretical efforts have been expended on prediction and cross-ecosystem comparison of particular patterns. It is now important to systematically confront these diverse observations within each ecosystem, from metabolism and density-dependence to variability and trophic cascades, to provide rigorous foundations for our understanding of trophic dynamics.

Another issue is the possibility for patterns to have causes outside the studied ecosystem. External energy subsidies, e.g. influxes of organic matter, or a predation range coupling multiple local communities, may be responsible for stronger trophic cascades [75] and top-heavy distributions [14, 76]. While a single pattern cannot rule out external causes, multiple stability and functioning patterns all pointing toward the same food chain structure would be a strong signal against this possibility.

This synthesis has been called for in conceptual frameworks [29], but it must become quantitatively precise if it is to solve long-standing empirical paradoxes. The relationships that we have summarized here should become part of a larger quantitative toolbox designed to provide better insight into the essential trophic structure of an ecosystem. For these predictions to hold in diverse ecological communities, they must be robust to the addition of complex structure, a question which emerging theoretical tools [77] and better integration with data may help answer in future studies.

## DATA AVAILABILITY

The metabolic and interaction data used for this study have been compiled with permission and deposited on figshare with the identifier: https://doi.org/10.6084/m9.figshare.6818288.v1. All code used for the analysis is available from the corresponding author upon request.

## ACKNOWLEDGMENTS

We thank J.-F. Arnoldi, N. Galiana Ibáñez, A. Sentis, D. Shanafelt and Y. Zelnik for comments during the elaboration of the manuscript. Our gratitude goes to A. Makarieva, C. White and their respective coauthors and sources for the data they compiled or collected, as well as all scientists who generously shared their work on the open GloBI and FishBase databases. This work was supported by the TULIP Laboratory of Excellence (ANR-10-LABX-41) and by the BIOSTASES Advanced Grant, funded by the European Research Council under the European Union’s Horizon 2020 research and innovation programme (666971).

